# N-Acyl Dopamines Induce Apoptosis in PC12 Cell Line via the O-1918-Sensitive non-CB1-non-CB2 Receptor

**DOI:** 10.1101/075796

**Authors:** Mikhail G. Akimov, Alina M. Ashba, Natalia M. Gretskaya, Vladimir V. Bezuglov

**Affiliations:** Shemyakin-Ovchinnikov Institute of Bioorganic Chemistry, Russian Academy of Sciences, Moscow, Russia

**Author notes:** Equally contributed authors (Mikhail G. Akimov, Alina M. Ashba). Correspondence to: Mikhail G. Akimov, Shemyakin-Ovchinnikov Institute of Bioorganic Chemistry, Russian Academy of Sciences, 117997, ul. Miklukho-Maklaya, 16/10, Moscow,Russia.

**Keywords:** N-acyl dopamine, cancer, cytotoxicity, apoptosis, non-CB1-non-CB2 receptor

## Abstract

Dopamine amides of long chain fatty acids (NADA) are a family of endogenous mammalian lipids with an unknown function; they are anti-proliferative for many cancer cell lines. The aim of this study was to identify the NADA receptor responsible for cell death induction. Using PC12 cells treated with NADA in combination with various receptor blockers, or pre-treated with the known receptor agonists to down-regulate the proposed receptor, NADA were shown to induce apoptosis. This activity was blocked only by the non-CB1-non-CB2 receptor antagonist O-1918, and by the pre-treatment with the agonists of this receptor ethanolamides of palmitic and oleic acids.

## Introduction

N-acyl dopamines (NADA) are biogenic conjugates of dopamine with long chain fatty acids. Their endogenous nature was first proposed independently by Bezuglov *et al.* (1) and Pokorski and Matysiak (2). Subsequently, NADA with arachidonic, docosahexaenoic, oleic, palmitic and stearic acid residues were found in the nervous system of *Rattus norvegicus* and *Bos taurus* (0.1-6 pmol/g wet weight) (3, 4), in two species of the *Hydra* genus (5) and in human plasma (6). The targets of NADA are vanilloid receptor TRPV1 and cannabinoid receptor CB1, as well as several other proteins (7).

Previously, others and we showed the ability of NADA to induce cell death in various cancer cell lines with a half-lethal concentration in range 2-50 μM (8–10). This activity depends on the structure of the fatty acid residue and on the modification of dopamine moiety (9) and thus is specific. However, the exact molecular target of NADA leading to cell death induction is still unknown.

The aim of this study was to identify the cell death mode and the NADA receptor target in rat pheochromocytoma PC12 cell line.

## Materials and Methods

Rat PC12 cell line was a kind gift of Dr. V. Shram (IMG RAS, Russia). RPMI 1640 cell medium, Hank’s solution, HEPES, phosphate-buffered saline, streptomycin, penicillin, amphotericin B, 3-(4,5-dimethylthiazol-2-yl)-2,5-diphenyltetrazolium bromide (MTT), and trypsin-EDTA solution were from PanEco, Moscow, Russia. D-glucose was from Sigma-Aldrich, USA. Fetal bovine serum was from PAA Laboratories, USA.

All test compounds were synthesized in the Laboratory of Oxylipins of Shemyakin-Ovchinnikov Institure of Bioorganic Chemistry, RAS, Russia as previously described (1) and stored only as ethanol or dimethyl sulfoxyde (DMSO) stocks under argon atmosphere at − 52°C.

Receptor blockers SR141716A (CB1), SR144528 (CB2), capsazepine (TRPV1), haloperidol (D2, D3), T0070907 (PPAR-gamma) and autophagy inhibitor hydroxychloroquine sulphate were from Sigma-Aldrich, USA; O-1918 (non-CB1-non-CB2) was from Cayman Europe, Estonia; pancaspase inhibitor Z-VAD-FMK and caspase fluorogenic substrates Ac-LEHD-AFC and Ac-DEVD-AFC were from Tocris Bioscience, UK.

PC12 cells were maintained at 37°C under 5% CO_2_ in the RPMI 1640 medium, supplemented with 2 mM L-glutamine, 7.5% fetal bovine serum, 100 U/ml penicillin, 0.1 mg/ml streptomycin and 0.25 μg/ml amphotericin B. Cells were passaged each 3 days and continiously grown for no more than 40 passages. Attached cells were detached with 0.25% trypsin in 0.53 mM EDTA in Hanks' salts. Cells were counted using a glass hemocytometer.

For analysis of the toxicity of NADA, the cells were plated in 96-well plates at a density of 1.25 × 10^4^ cells per well and grown for 3 days. The dilutions of test compounds prepared in DMSO and dissolved in the conditioned culture medium were added to the cells in triplicate for each concentration and incubated for 18 h; the inhibitors were added 1 h before N-acyl dopamines. The incubation time was chosen based on the most pronounced differences between the compounds tested. The final DMSO concentration was 0.5%. Negative control cells were treated with 0.5% DMSO. Positive control cells were treated with 3.6 μl of 50% Triton X-100 in ethanol per 200 μl of cell culture medium. Separate controls were without DMSO (no difference with the control 0.5% DMSO was found, data not shown). The effect of test substances on the cell viability was evaluated using the MTT test (based on the MTT dye reduction by mitochondria of living cells) (11).

For the MTT test, after removal of the medium with the test compounds, the cells and the controls were incubated for 1.5 h with 0.5 mg/ml of MTT in Hank's solution, supplemented with 10 mM of D-glucose. After this incubation, the solution was removed and the cells were dissolved in DMSO. The amount of the reduced dye was determined colorimetrically at 594 nm with a reference wavelength 620 nm using an EFOS 9505 photometer (Sapphire, Moscow, Russia). Additionally, the attachment and cell shape of adherent cells were evaluated microscopically.

To perform a pharmacological downregulation of the surface receptor sensitive to O-1819 and on-CB1-non-CB2 agonists, the cells were incubated with these substances for 3 days. At the end of the incubation, the cells were washed with the fresh medium and NADA solution was added as described before for 1 h, after which it was replaced with a fresh cultivation medium for 24 h. Assessment of the cell viability was carried out using the MTT assay.

The determination of caspase activity was performed using the specific substrates labeled with a fluorescent label AFC. The experiment was performed in 6-well plates. After the 24 hour incubation, the medium was discarded and 1 ml of the lysis buffer (20 mM HEPES, 10 mM KCl, 1.5 M MgCl_2_, 0.5% NP-40, 1 mM dithiothreitol, 1 mg/ml pepstatin A, 1 mg/ml aprotinin, 5 mg/ml leupeptin, 1 mM PMSF) was added to the wells and left for 10 minutes. Then the cells were resuspended, the cell lysate was centrifuged using a microcentrifuge at 15,000 x g for 30 min, and the supernatant was collected. Activity determination of caspase-3 and −9 was performed by mixing the supernataint aliquote and the corresponding specific substrate in the reaction buffer (50 mM HEPES, 10% sucrose, 0.1% CHAPS, 100 mM dithiothreitol) at the ratio of 1:1. Thereafter, the samples were incubated for 30 min at 37°C. The following substrates were used for caspases: for caspase-3, Ac-DEVD-AFC (50 uM) (Ac-Asp-Glu-Val-Asp-AFC), for caspase-9, Ac-LEHD-AFC (50 μM) (Ac - Leu-Glu-His-Asp-AFC). A pan-caspase inhibitor Z-VAD-FMK (50 μM) was used as a negative control, and staurosporine (10 mM) as the positive one. The determination of the liberated AFC was performed using a plate fluorimeter (Hidex, Finland), λ_ex_ = 505 nm, λ_em_ = 400 nm.

To confirm the induction of apoptosis, an apoptosis detection kit based on the fluorescent dye Apo-TRACE (Sigma-Aldrich, USA), was used as recommended by the manufacturer. The method principle is the selective accumulation of the dye in the cells undergoing apoptosis. The fluorescence intensity was measured using a plate fluorimeter (Hidex, Finland), λ_ex_ = 350 nm, λ_em_ = 420 nm.

To check the apoptosis-specific DNA fragmentation, DNA was extracted using the Biotechrabbit Blood DNA kit according to the manufacturer's instructions and analyzed using agarose gel electrophoresis (gel percentage 2.0, constant voltage 137 V, Tris-Borate-EDTA buffer).

Each experiment was repeated three times. Curves for calculating the 50% lethal dose (LD_50_) were generated using the GraphPad Prism software (www.graphpad.com). Data are presented as mean ± standard deviation. Data were compared using the unpaired Student's *t* test; *p* values of 0.05 or less were considered significant.

## Results

In our previous work, we found dopamine amides of arachidonic (AA-DA), docosahexaenoic (DHA-DA) and oleic (Ol-DA) acids to be cytotoxic for PC12 cells with LC_50_=2-4 μM (10). In this work, we focused on the evaluation of the cell death type, induced by these substances in PC12 cells, and on the identification of the surface receptor engaged in this process.

The minimum time of NADA exposition, enough for cell death induction was shown to be 1 h; even if the substance was washed out, the cell death program unrolled from this time point (data not illustrated). The pre-treatment with an autophagy inhibitor hydroxychloroquine (12) did not alter cell survival, while the pan-caspase inhibitor Z-VAD-FMK was able to prevent NADA cytotoxicity (Figure 1). From this we suggested that NADA induce apoptosis, and confirmed it using three additional methods: fluorometric caspase activation assay (Figure 2A), ApoTRACE apoptosis detection kit (Figure 2B), and DNA laddering assay (Figure 2C). We observed activation of both caspase 3 and 9, and, together with our previous data on the ability of NADA to induce ROS accumulation (10), we concluded that these substances induce apoptosis via the intrinsic pathway.

**Figure 1.**
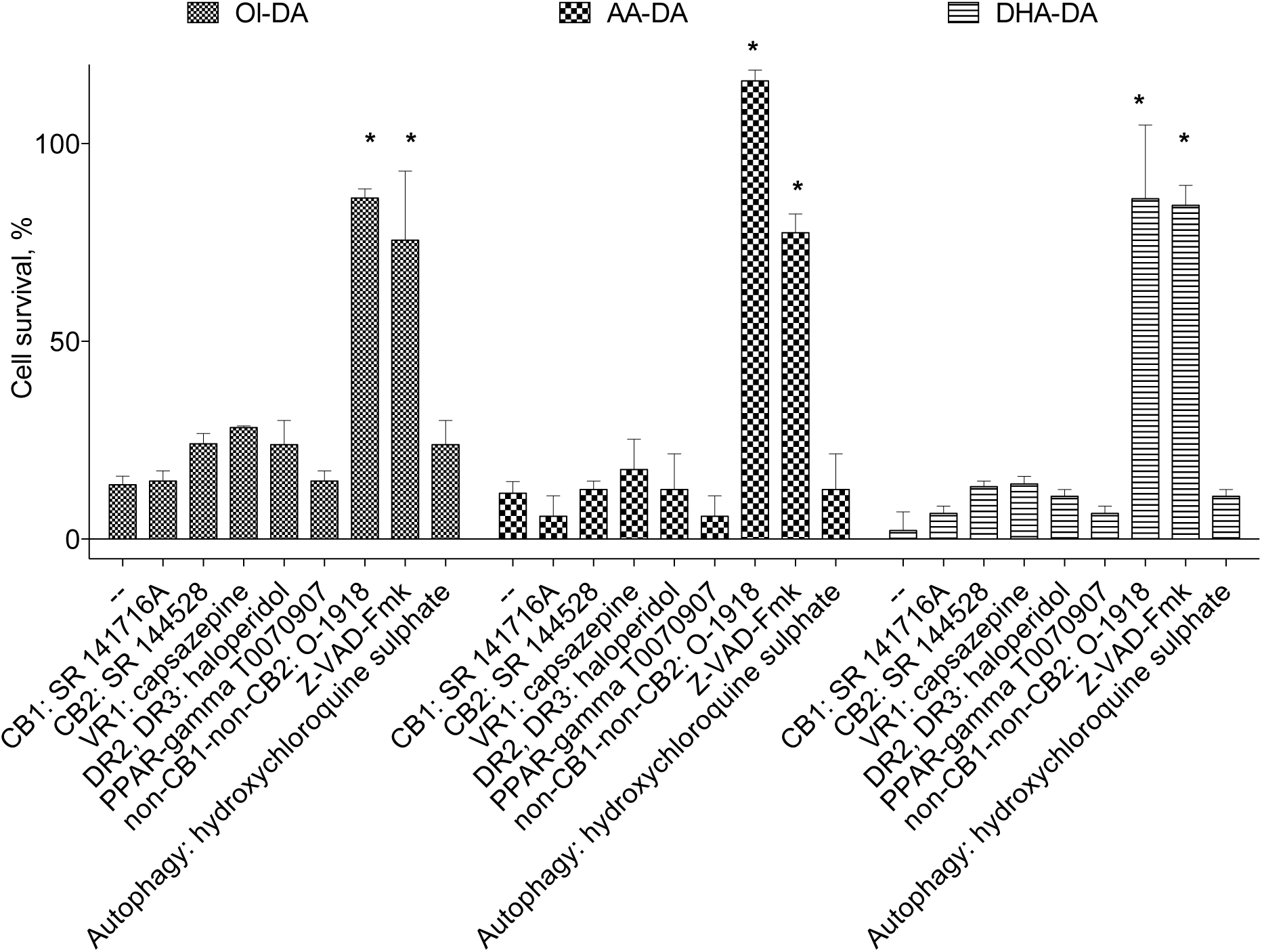
Receptor blocker sensitivity of the cytotoxic effect of dopamine amides of arachidonic (AA-DA), docosahexaenoic (DHA-DA) and oleic (Ol-DA) acids. Simultaneous addition; NADA, 3 μM, SR 141716A, 0.5 μM, SR 144528, 0.5 μM, capsazepine, 2 μM, haloperidol, 10 μM, T0070907, 1 μM, Z-VAD-Fmk, 50 μM, hydroxychloroquine sulphate, 10 μM, O-1918, 3 μM. MTT test data for 18 h incubation time, mean±standard deviation; * statistically significant difference from NADA without blockers, *p*<0.05 (multiple ANOVA with the Tukey post test)

**Figure 2.**
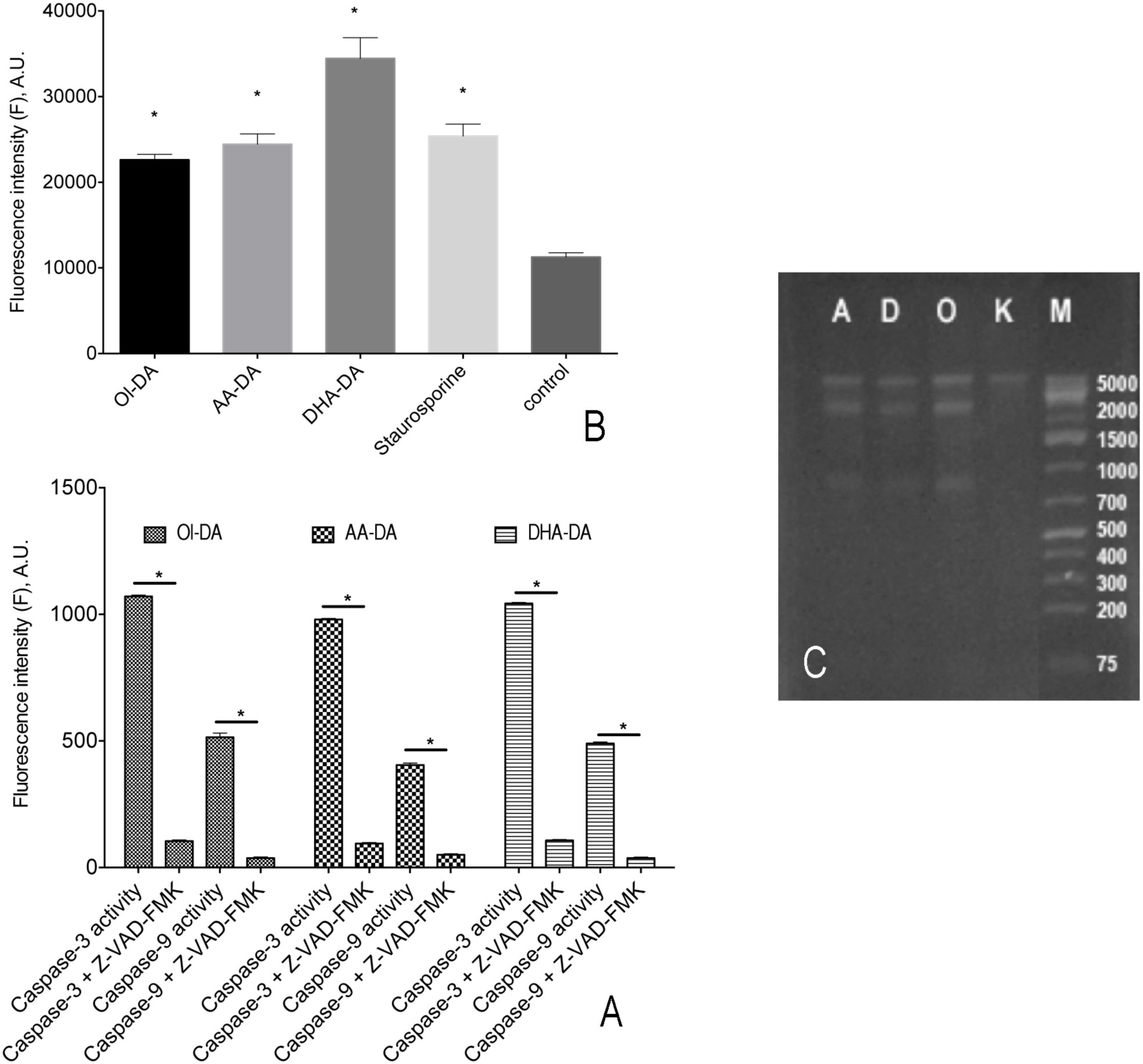
Apoptosis induction by dopamine amides of arachidonic (AA-DA), docosahexaenoic (DHA-DA) and oleic (Ol-DA) acids in the PC12 cells. A, Caspase activation. Incubation time with NADA was 5 h. Fluorescence data for caspase 3 substrate Ac-DEVD-AFC and caspase 9 substrate Ac-LEHD-AFC cleavage; mean±standard deviation; * statistically significant difference from the control with Z-VAD-Fmk, p<0.05 (Student's ttest). B, ApoTRACE accumulation. Fluorescence data for 5 h incubation time, mean±standard deviation; * statistically significant difference from control, p<0.001 (multiple ANOVA with the Dunnett post test). C, DNA laddering. Agarose gel electrophoresis for 5 h incubation with ethidium bromide staining, A, AA-DA, D, DHA-DA, O, Ol-DA, K, control, M, markers

To identify the receptor responsible for NADA cell death induction, we pre-incubated the cells with a panel of blockers of putative and known endocannabinoid and endovanilloid receptors (Figure 1). The only active substance was O-1918, a non-CB1-non-CB2 receptor blocker (13). To further confirm the participation of an O-1918-sensitive receptor in NADA cytotoxicity, we performed a pre-incubation with the inhibitor for 72 h, as this technique is known to diminish surface receptor density (14). Then the inhibitor was washed out, and the cells were incubated with NADA for 1 h with washing out of the substances at the end. We found that such pretreatment led to a significant decrease of NADA cytotoxicity (Figure 3), which indicates the change of the receptor density or sensitivity.

The non-CB1-non-CB2 receptor group comprises a number of proteins, which include as-yet-unidentified/putative receptor(s), and established orphan GPCRs, namely GPR92, GPR55, and GPR119 (15). To further evaluate the participation of these receptors in the NADA cell death induction, we used four known ligands of the receptors of this group, namely the GPR55 ligand N-palmitoyl ethanolamide (Palm-EA (16)), the GPR55 (16) and GPR119 (17) ligand N-oleoyl ethanoleamide (Ol-EA), the GPR92 ligand N-arachidonoyl glycine (AA-Gly (18)), and the putative endothelial cannabinoid receptor ligand N-arachidonoyl serine (AA-Ser (19)). First, we checked the ability of these substances to mimic NADA action on PC12 cells and found that they indeed induced cell death, which was blocked by O-1918 (Figure 4). Next we incubated the cells with a non-lethal concentration of each of these substances for 72 h to induce the appropriate receptor downregulation and tested the activity of NADA on the survived cells. Palm-EA and Ol-EA pretreatment was able to almost completely prevent the NADA cytotoxicity, while AA-Gly and AA-Ser were without effect (Figure 5).

**Figure 3.**
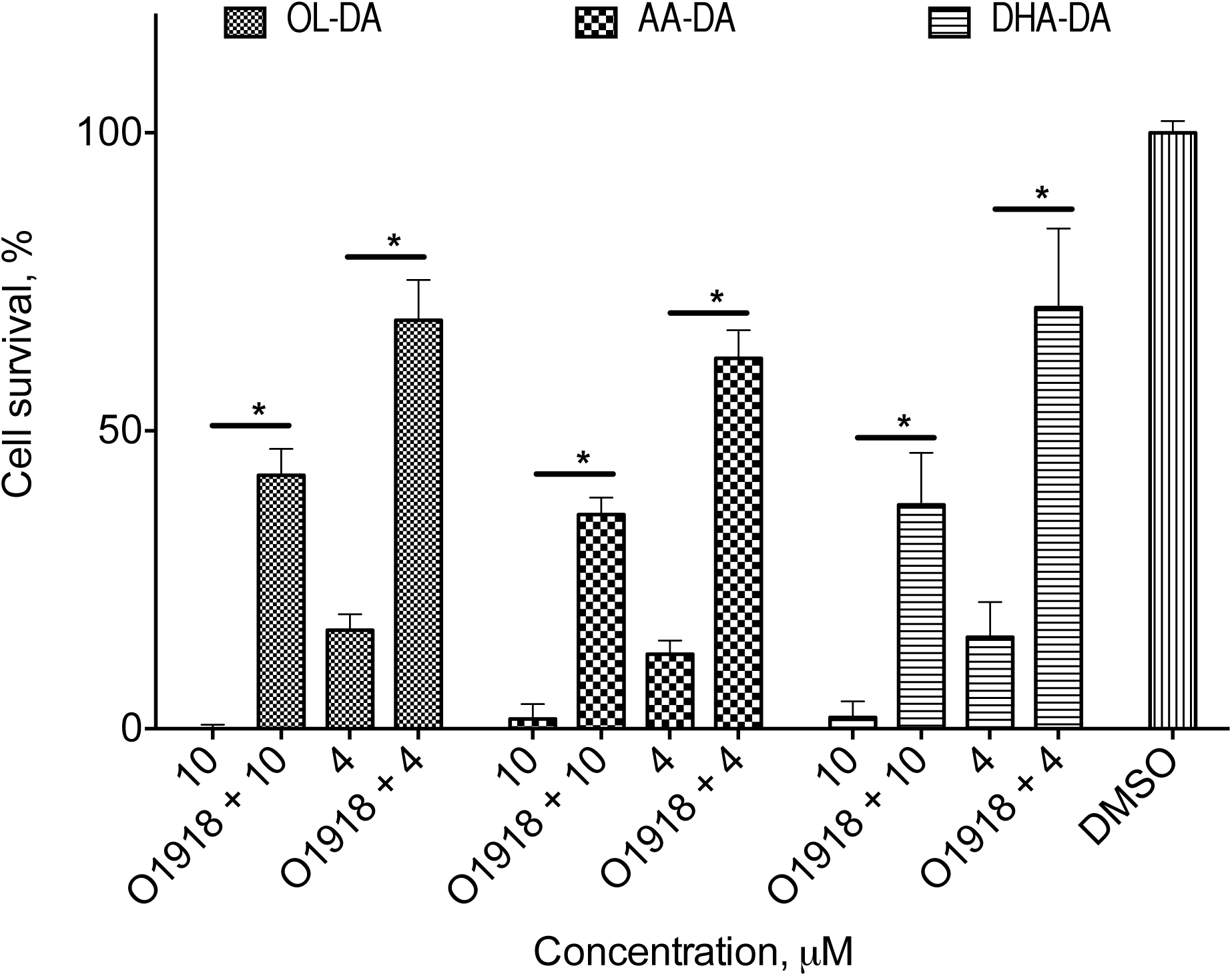
Cytotoxicity of dopamine amides of arachidonic (AA-DA), docosahexaenoic (DHA-DA) and oleic (Ol-DA) after the 72 h O-1918 pre-treatment and washing-out of the PC12 cells. MTT test data for 18 h incubation time, mean±standard deviation; * statistically significant difference, *p*<0.05 (multiple ANOVA with the Tukey post test)

**Figure 4.**
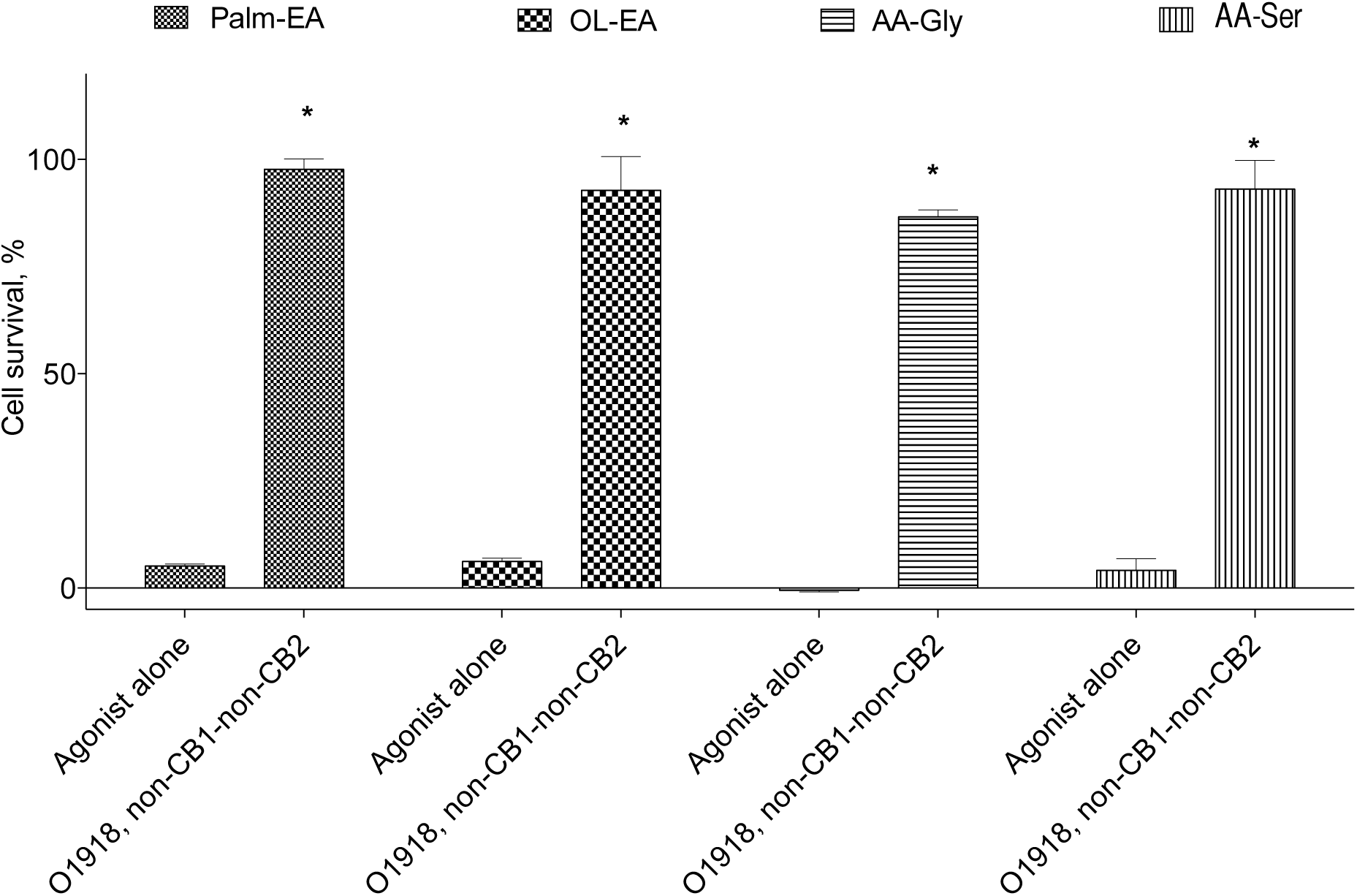
Cytotoxicity and inhibitor sensitivity of the known non-CB1-non-CB2 receptor agonists for the PC12 cells. Concentration of the agonists 25 μM. MTT test data for 18 h incubation time, mean±standard deviation; * statistically significant difference from the appropriate agonist alone, *p*<0.05 (multiple ANOVA with the Tukey post test)

**Figure 5.**
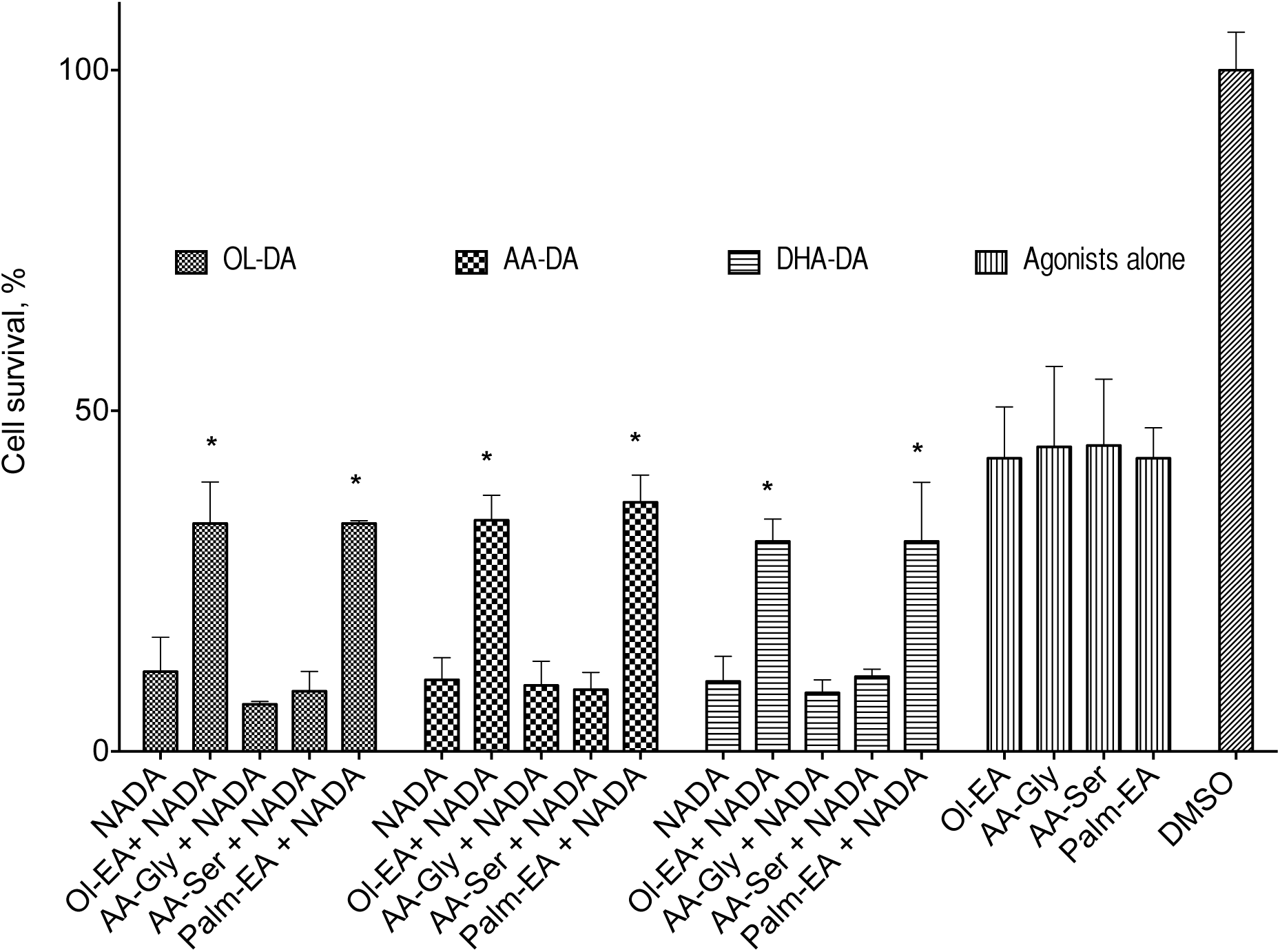
Cytotoxicity of dopamine amides of arachidonic (AA-DA), docosahexaenoic (DHA-DA) and oleic (Ol-DA) after the 72 h non-CB1-non-CB2 agonists pre-treatment and washing-out of the PC12 cells. The concentration of the agonists (Palm-EA, 15 µM, Ol-EA, 5 µM, AA-Gly, 10 µM, AA-Ser, 5 µM) was chosen to kill 50% of the cells, and the concentration of the dopamine amides (5 µM)-to kill 80% of the cells. MTT test data for 18 h incubation time, mean±standard deviation; * statistically significant difference from the appropriate dopamine amide alone, *p*<0.05 (multiple ANOVA with the Tukey post test)

## Discussion

The main result of this research is the demonstration for the first time that NADA induce apoptosis of a cancer cell line via the activation of the nonclassical cannabinoid non-CB1-non-CB2 receptor. The cytotoxic properties of NADA and their analogs was described in literature many times (8–10), with some insights for the intracellular ROS– and NO-dependent mechanism of their action (10). However, our study is the first one to identify the cell surface molecular target of these effects. We checked a set of major cellular receptor inhibitors that blocked activity of surface classical cannabinoid (CB1, CB2), dopamine (D2, D3), vanilloid (TRPV1), non-CB1-non-CB2 receptors, as well as an inhibitor of nuclear receptor PPAR_γ_, known to be involed in pro-apoptotic action of capsaicin – structural analog of NADA (20). From this set of inhibitors, only O-1918 rescued cells from the toxic action of NADA; all other inhibitors were ineffective (Fig. 1). O-1918, which lacks detectable affinity for CB1 and CB2 receptors, is a structural analog of abnormal cannabidiol and was reported to be a neutral antagonist of the putative endothelial non-cannabinoid receptor (21, 22). The critical involvement of the non-CB1-non-CB2 receptor in the cytotoxic effect of NADA is supported by the results of pharmacological blocade of the aforementioned receptor using a prolonged incubation with O-1918, Palm-EA and Ol-EA, which dramatically increased viability of the PC12 cells upon NADA treatment (Fig. 3, 5). Activation of this putative receptor with three tested NADA (Ol-DA, AA-DA or DHA-DA) leads to the promotion of apoptotic death of PC12 cells, which was confirmed by three independent tests: caspase activation, DNA laddering and ApoTRACE assays. We can speculate that the downstream mechanism of NADA-induced apoptosis initiated by interaction with putative non-CB1-non-CB2 receptor is linked to the reactive oxygen species overproduction, as was recently demonstrated by us (10).

O-1918 can block several non-cannabinoid receptors (glial, GPR55, GPR18) (22). The NADA cytotoxicity change after the non-CB1-non-CB2 agonists pretreatment could possibly narrow this list to GPR55, which is in line with the presence of the functional GPR55 in the PC12 cells (23) and with the ability of the endocannabinoid anandamide to induce cholangiocarcinoma cell death via the GPR55 activation (24). Another indirect evidence for the GPR55 participation is that this receptor is able to switch its signal transduction pathways depending on the ligand (25), and that anandamide activation of this receptor induced cell death (24), while the activation by lysophosphatidylinositol did not (23). However, the differentiation of the observed NADA effects between all these receptors, as well as further elucidation of the intracellular transduction pathways during the NADA-induced apoptosis of cancer cells were not the subject of this paper and will be reported elsewhere.

## Acknowledgements

The work was partially supported by the grant of Russian President MK-3842.2015.4 and grant of Russian Foundation for Basic Research # 16-04-00729a.

## References

1. Bezuglov VV, Manevich Y, Archakov AV, Bobrov MYu, Kuklev DV, Petrukhina GN, et al. Artificially functionalized polyenoic fatty acids as new lipid bioregulators. Russ J Bioorg Chem. 1997;23(3):211–20.

2. Pokorski M, Matysiak Z. Fatty acid acylation of dopamine in the carotid body. Med Hypotheses. 1998;50(2):131–3.

3. Huang SM, Bisogno T, Trevisani M, Al-Hayani A, De Petrocellis L, Fezza F, et al. An endogenous capsaicin-like substance with high potency at recombinant and native vanilloid VR1 receptors. Proc Natl Acad Sci U S A. 2002;99(12):8400–5.

4. Chu CJ, Huang SM, De Petrocellis L, Bisogno T, Ewing SA, Miller JD, et al. N-oleoyldopamine, a novel endogenous capsaicin-like lipid that produces hyperalgesia. J Biol Chem. 2003;278(16):13633–9.

5. Ostroumova TV, Markova LN, Akimov MG, Gretskaia NM, Bezuglov VV. Docosahexaenoyl dopamine in freshwater hydra: effects on regeneration and metabolic changes. Russ J Dev Biol. 2010;41(3):164–7.

6. Hauer D, Schelling G, Gola H, Campolongo P, Morath J, Roozendaal B, et al. Plasma concentrations of endocannabinoids and related primary fatty acid amides in patients with post-traumatic stress disorder. PLoS One. 2013;8(5):e62741.

7. Akimov MG, Bezuglov VV. N-Acylated Dopamines: A New Life for the Old Dopamine. In: Kudo E, Fujii Y, editors. Dopamine: Functions, Regulation and Health Effects. NY: Nova Science Publishers; 2012. p. 49–80.

8. Visnyei K, Onodera H, Damoiseaux R, Saigusa K, Petrosyan S, De Vries D, et al. A molecular screening approach to identify and characterize inhibitors of glioblastoma multiforme stem cells. Mol Cancer Ther. 2011.

9. Akimov MG, Gretskaya NM, Zinchenko GN, Bezuglov VV. Cytotoxicity of endogenous lipids N-acyl dopamines and their possible metabolic derivatives for human cancer cell lines of different histological origin. Anticancer Res. 2015;35(5):2657–61.

10. Ashba AM, Akimov MG, Gretskaya NM, Bezuglov VV. N-Acyl Dopamines Induce Cell Death in PC12 Cell Line via Induction of Nitric Oxide Generation and Oxidative Stress. Doklady Biochemistry and Biophysics,. 2016;467:1–4.

11. Mosmann T. Rapid colorimetric assay for cellular growth and survival: application to proliferation and cytotoxicity assays. J Immunol Methods. 1983;65(1-2):55–63.

12. Lee H-O, Mustafa A, Hudes GR, Kruger WD. Hydroxychloroquine Destabilizes Phospho-S6 in Human Renal Carcinoma Cells. PLoS One. 2015;10(7):e0131464.

13. Offertaler L, Mo F-M, Batkai S, Liu J, Begg M, Razdan RK, et al. Selective ligands and cellular effectors of a G protein-coupled endothelial cannabinoid receptor. Mol Pharmacol. 2003;63(3):699–705.

14. Law PY, Hom DS, Loh HH. Opiate receptor down-regulation and desensitization in neuroblastoma X glioma NG108-15 hybrid cells are two separate cellular adaptation processes. Mol Pharmacol. 1983;24(3):413–24.

15. Godlewski G, Kunos G. Overview of nonclassical cannabinoid receptors. endoCANNABINOIDS: Springer; 2013. p. 3–27.

16. Ryberg E, Larsson N, Sjogren S, Hjorth S, Hermansson NO, Leonova J, et al. The orphan receptor GPR55 is a novel cannabinoid receptor. Br J Pharmacol. 2007;152(7):1092–101.

17. Godlewski G, Offertaler L, Wagner JA, Kunos G. Receptors for acylethanolamides-GPR55 and GPR119. Prostaglandins Other Lipid Mediat. 2009;89(3-4):105–11.

18. Oh DY, Yoon JM, Moon MJ, Hwang JI, Choe H, Lee JY, et al. Identification of farnesyl pyrophosphate and N-arachidonylglycine as endogenous ligands for GPR92. J Biol Chem. 2008;283(30):21054–64.

19. Milman G, Maor Y, Abu-Lafi S, Horowitz M, Gallily R, Batkai S, et al. N-arachidonoyl L-serine, an endocannabinoid-like brain constituent with vasodilatory properties. Proc Natl Acad Sci U S A. 2006;103(7):2428–33.

20. Kim CS, Park WH, Park JY, Kang JH, Kim MO, Kawada T, et al. Capsaicin, a spicy component of hot pepper, induces apoptosis by activation of the peroxisome proliferator-activated receptor gamma in HT-29 human colon cancer cells. Journal of medicinal food. 2004;7(3):267–73.

21. Pertwee RG, Howlett AC, Abood ME, Alexander SP, Di Marzo V, Elphick MR, et al. International Union of Basic and Clinical Pharmacology. LXXIX.Cannabinoid receptors and their ligands: beyond CB(1) and CB(2). Pharmacological reviews. 2010;62(4):588–631.

22. Abood ME, Sorensen RG, Stella N. Endocannabinoids : actions at non-CB1. New York: Springer; 2013.

23. Obara Y, Ueno S, Yanagihata Y, Nakahata N. Lysophosphatidylinositol causes neurite retraction via GPR55, G13 and RhoA in PC12 cells. PLoS One. 2011;6(8):e24284.

24. Huang L, Ramirez JC, Frampton GA, Golden LE, Quinn MA, Pae HY, et al. Anandamide exerts its antiproliferative actions on cholangiocarcinoma by activation of the GPR55 receptor. Lab Invest. 2011;91(7):1007–17.

25. Henstridge CM, Balenga NA, Schroder R, Kargl JK, Platzer W, Martini L, et al. GPR55 ligands promote receptor coupling to multiple signalling pathways. Br J Pharmacol. 2010;160(3):604–14.

